# A novel chemogenomic screening platform for scalable antimalarial drug target identification

**DOI:** 10.1101/2024.08.27.609875

**Authors:** Christopher Bower-Lepts, Mukul Rawat, William Collins Keith, Gareth Girling, Madeline R Luth, Krypton Carolino, Katharina Boroviak, Rachael Coyle, Frank Schwach, Cindy Smidt, Ellen Bushell, Sonia Moliner-Cubel, Maria D Gomez-Lorenzo, Francisco-Javier Gamo, Elizabeth A. Winzeler, Oliver Billker, Marcus C.S. Lee, Julian C. Rayner

## Abstract

Large-scale chemical-genetic screening, or chemogenomics, can faciliate rapid and scalable drug target identification. To establish a chemogenomic screen for antimalarial drug target identification, we leveraged ∼600 *Plasmodium berghei* artificial chromosomes (PbACs) encoding potential drug targets to generate a systematic overexpression library. PbACs were engineered with DNA barcodes, enabling their quantification within mixed pools using next generation sequencing (barcode sequencing or BarSeq). Pooled transfection of PbACs into the highly genetically tractable *Plasmodium knowlesi* demonstrated efficient vector uptake and transcription of encoded *P. berghei* genes. Parasite pools were exposed to antimalarial candidates, with pilot screens probing for known gene-compound associations identifying their targets with high sensitivity. Screening antimalarial inhibitors with unknown mechanisms of action successfully identified *pi4k* as the target for one novel compound, which was subsequently validated using *in vitro* evolution in *Plasmodium falciparum* parasites. This sensitive and scalable chemogenomics platform therefore represents a valuable early-stage tool for antimalarial target identification.

## Introduction

Malaria exacts a devastating public health burden in endemic countries, with over 200 million cases and 600,000 deaths recorded in 2023, largely due to infections with *Plasmodium falciparum*^1^. Despite the combined efforts of vector control and drug prophylaxis, the therapeutic administration of antimalarial medicines remains central to reducing malaria mortality, yet has been repeatedly undermined by the evolution and spread of drug resistant parasites. Most recently, the emergence of resistance to artemisinin in Southeast Asia^2,3^ and East Africa^4,5^, threatens a devastating reversal of current malaria control. New drugs, particularly drugs employing novel mechanisms of action, are urgently needed to replenish the antimalarial pipeline.

To identify leads for new antimalarials, multiple large-scale phenotypic screens have been conducted to identify compounds with activity against *P. falciparum* asexual blood stage growth *in vitro*^6,7,8^. These screens have been highly successful, identifying thousands of potent scaffolds (<10 uM). Several candidates have progressed to clinical trials^9,10^, however, many more fail, and additional candidates are always required. Compound development is frequently stalled by a lack of a known molecular target or mechanism of action. Target identification can significantly facilitate medicinal chemistry optimization of activity, selectivity, and a compounds safety profile, increasing the likelihood of positive drug development outcomes^11^. Target deconvolution for antimalarials has largely centred on a few techniques, including *in vitro* evolution of resistance and whole genome analysis (IVIEWGA), thermal proteome profiling (TPP), chemoproteomics and metabolomics^12,13,14^. IVIEWGA has been the most widely and successfully employed target identification method, discovering numerous targets and augmenting the development of several clinical candidates^15,16,17^. However, the approach is time-consuming, involving long-term parasite culture under compound selection. In some cases, no clear target emerges or resistance cannot be generated, and it is difficult to scale to hundreds of compounds. Considering the many potential antimalarial therapies with unidentified targets, novel approaches for scalable target identification would significantly facilitate the development of phenotypic hits into drugs.

Large-scale chemical-genetic interaction screens, or chemogenomics, have been used to identify drug mechanisms-of-action in several disease models^18,19^. This typically involves the systematic alteration of expression levels, via under- or over-expression of targets at the transcript or protein level and exposure to compounds of interest. Mutant strains in which a given target’s expression level is reduced or increased are predicted to show sensitization or resistance, respectively, to compounds that interact with that target. Performing chemogenomic screens at scale therefore necessitates the generation of large numbers of transgenic lines. Although loss-of-function chemogenomic screens have been carried out using pools of random piggyBac transposon mutants^20, 21^ the low transfection efficiency of *P. falciparum* makes the targeted generation of many transgenic lines challenging. As a result, there have been no attempts at large-scale over-expression chemical-genetics in *Plasmodium* parasites to date.

To develop an over-expression chemogenomic tool for antimalarial drug target identification, we combined three key technological developments. Firstly, we used the zoonotic human pathogen *Plasmodium knowlesi*, which has a manifold higher transfection efficiency than *P. falciparum*^22,23^, permitting the transfection of large numbers of vectors in blood-stage human *Plasmodium* parasites for the first time. Secondly, we applied Barcode sequencing (Barseq), where next-generation sequencing (NGS) is used to quantify the growth of individual barcoded mutant strains grown in mixed pools, as has been previously employed for genome-scale knockout screening of the *Plasmodium berghei* genome^24,25^. Finally, we leveraged a library of *Plasmodium* artificial chromosomes which provides a resource to drive gene overexpression through addition of extra copies of genes of interest (Billker Lab, Unpublished Data). While this library was developed for the rodent model *P. berghei* (*P. berghei* artificial chromosomes or PbACs) (Figure 1A), promoter and terminator elements from one *Plasmodium* species are routinely used to drive transgene expression in other *Plasmodium* species in plasmid-based approaches, rendering it possible that PbACs could be utilised to generate a library of overexpression strains in *P. knowlesi*.

**Figure 1.**
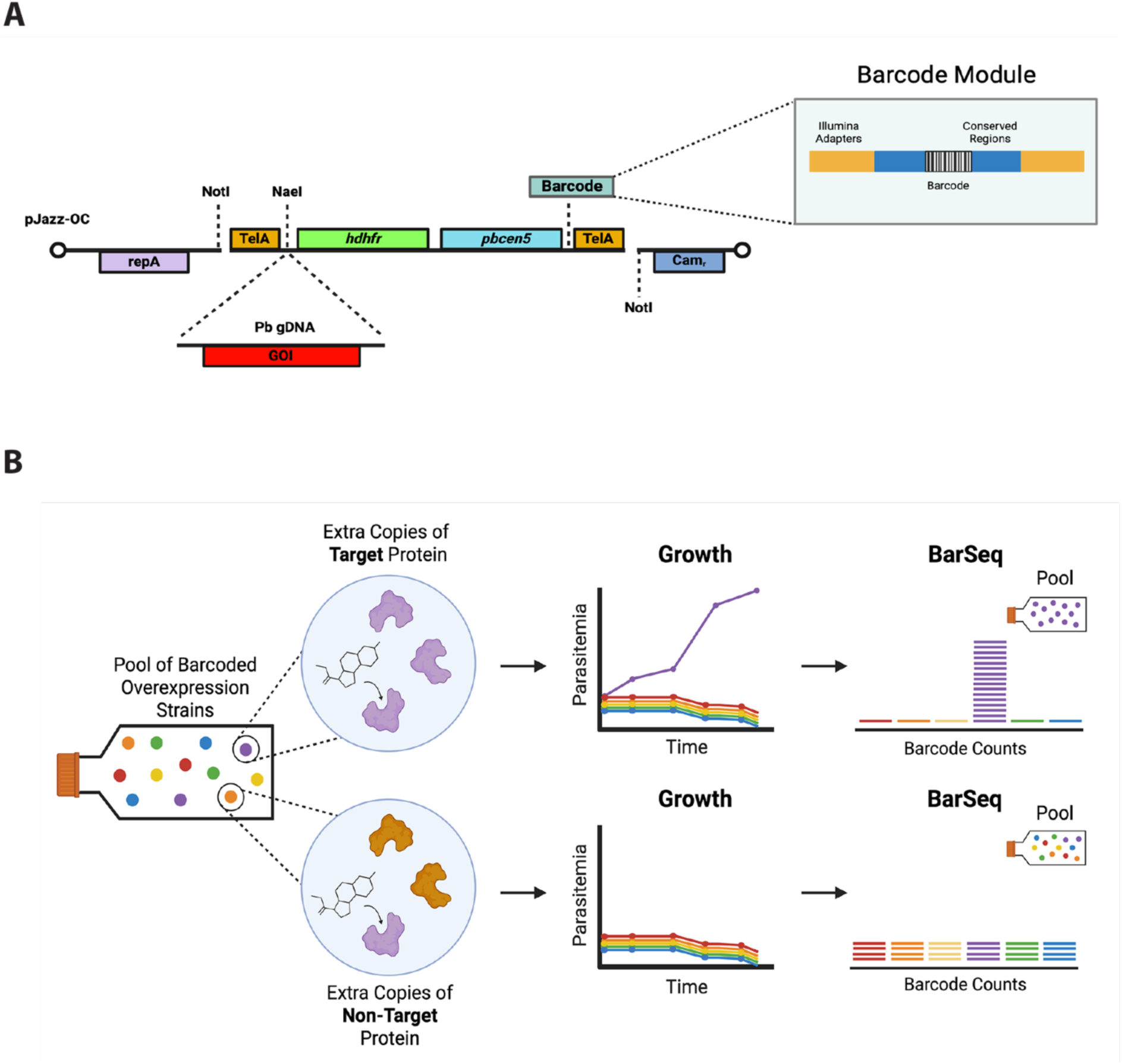
Establishing an *in vitro* chemogenomic screening approach in *Plasmodium* parasites. (**A**) Representative schematic diagram of the *Plasmodium berghei* artificial chromosome (PbAC) construct with barcode molecule added by recombineering. Key features include a centromere (*pbcen5*), telomeres (TelA) and human dihydrofolate resistance cassette encoding resistance to pyrimethamine (*hdhfr*) for drug selection. (**B**) Schematic overview of chemogenomic screening approach for target identification. Pools of barcoded overexpression lines are selected with antimalarial inhibitors. Mutants which are expressing extra copies of the compounds target genes may possess a selective advantage in pools relative to strains overexpressing non-target genes. Resistant strains grow to higher proportions in pools, while susceptible strains decrease in proportion. BarSeq on the pre-and post-selection populations demonstrates enrichment of barcodes for the gene encoding the target protein.

In this manuscript we combine these tools by introducing unique DNA barcodes into a subset of PbACs that contain potential drug targets, transfecting pools of barcoded PbACs into *P. knowlesi,* selecting pools in the presence of antimalarial inhibitors, and using BarSeq to identify genes whose over-expression conferred a growth advantage in the presence of the compounds (Figure 1B). This approach confirmed the known target for three test antimalarial compounds, and identified the target for a novel compound which was then confirmed in *P. falciparum* using IVIEWGA. This novel chemogenomic approach has significant potential to scale up the critical task of antimalarial drug target identification.

## Results

### Generation of a barcoded PbAC library for chemogenomics

To adapt the PbAC library for chemogenomic screening, we introduced unique DNA barcodes into individual PbACs, enabling the employment of BarSeq to demultiplex and quantify individual vectors in complex pools (Figure 1A). To avoid investing effort into unlikely targets and to limit the complexity of parasite pools when screening, only a subset of the PbAC library was prioritised for barcoding. Genes encoding likely drug targets (enzymes, transporters and membrane proteins that were likely to be non-dispensible for blood-stage growth, see methods for further details) and with favourable characteristics for successful target expression (PbAC includes promoter region ≥ 1 kb, terminator region ≥ 0.5 kb), were selected for barcoding (Figure S1). Recombinase-mediated engineering (recombineering) was used to engineer unique barcodes into individual PbAC constructs, utilising homologous recombination in *E. coli* mediated by bacteriophage lambda *redγβα* recombinase enzymes^26^, and recovered clones were end-sequenced to confirm successful barcode incorporation. Recombineering was attempted on 730 PbACs, yielding 585 unique barcoded constructs, covering 589 *P. berghei* genes (Table S1). The majority of barcoded PbACs contained the entire ORF and UTRs of a single gene (a small minority contained a maximum of two), encompassing a total construct size of 20-23 kb. Among the target genes in the final library were over 20 previously-validated antimalarial drug targets^12^, including several well-characterised *Plasmodium* targets which could serve as positive controls in target identification assays (Table S1).

### PbACs can be used for systematic target overexpression in *P. knowlesi*

*P. knowlesi* has previously demonstrated a high capacity for the parallel uptake of multiple vectors when transfected with pools of plasmids^23^, but this has not yet been established with large, linear constructs such as PbACs. To explore this, *P. knowlesi* A1H1 parasites were transfected with PbAC pools of increasing size, and BarSeq was performed pre- and post-transfection to determine the complexity of PbAC uptake. *P. knowlesi* demonstrated a high capacity for PbAC uptake in smaller pools of up to 29 PbACs (Figure S2). To test the limits of population complexity that could be established in a single transfection event, the input pool size was increased to 96 PbACs; all input vectors were successfully detected in the transfected population (Figure 2A). These results indicated an efficiency of PbAC uptake by *P. knowlesi* similar to that observed with small plasmids^23^, and establishing that highly complex parasite pools can be generated in a single transfection.

**Figure 2.**
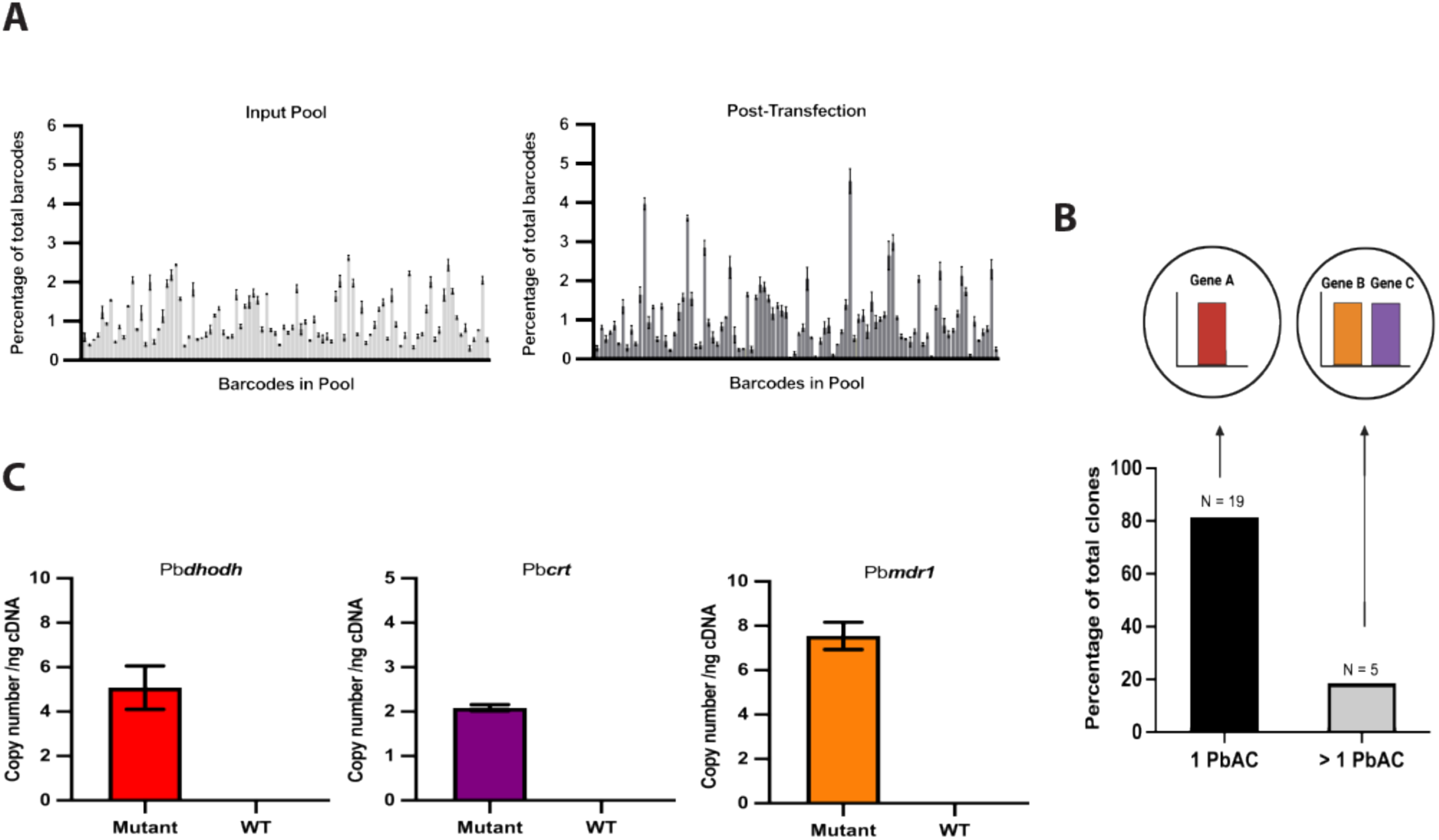
Validating PbACs as a vector to generate an overexpression library in *Plasmodium*. (**A**) BarSeq on the pre-transfection (Input Pool) and post-transfection population (Post-Transfection) for *P. knowlesi* parasites transfected with a pool of 96 PbAC vectors. Each bar denotes the proportion of total barcodes represented by a given PbAC barcode in the pool, corresponding to an individual construct in the library. Error bars represent standard deviation (SD). (**B**) The proportion of transfectant lines obtained by limited dilution cloning that contained one or multiple PbAC barcodes, as determined using BarSeq. Numbers above bars represent the number of clones identified in each case (total N = 24). (**C**) Transcription of *P. berghei* target genes encoded by PbACs measured in clonal parasite populations and *P. knowlesi* WT parasites using RT-qPCR, assayed in triplicate. Error bars represent SD.

To establish whether this complexity was accounted for by many parasites taking up one PbAC or individual parasites taking up multiple PbACs, clones were derived from transfectant pools using limiting dilution cloning and BarSeq used to determine how many different PbAC barcodes could be detected within each clone. In most clones only a single PbAC barcode was identified, indicating that primarily only one PbAC is taken up during transfection (Figure 2B) and that parasites will overexpress a single target, ideal for interrogating single gene-drug relationships. In rare cases, a maximum of two different PbACs could be detected in a single clone (Figure 2B & Figure S3), potentially reflecting the uptake of two PbACs following transfection, although mixed clones may also account for some of this phenomenon.

Finally, to establish that PbAC-encoded *P. berghei* genes were successfully transcribed by the *P. knowlesi* transcription machinery, RT-qPCR was used to detect *P. berghei* gene transcripts in PbAC transfectants (Figure 2C). Transfection of single PbACs was used to generate clonal lines, from which cDNA was assayed to measure transcript levels of the known drug targets *dhodh, crt* and *mdr1*. All three lines had confirmed transcription of these *P. berghei* genes, whereas no expression of these genes was detected in parental wildtype (WT) parasites (Figure 2C), or from negative control templates derived from PbAC clonal lines that were not subjected to reverse transcription ( -RT), ensuring that amplification was not as a result of contaminating gDNA. This indicates successful transcription of *P. berghei* genes, confirming that PbACs could be used to drive target overexpression in *P. knowlesi*.

### Testing the chemogenomic approach using pilot compounds with known modes of action

Having established that pooled transfection of barcoded PbACs could generate complex populations of overexpression strains in *P. knowlesi*, we next tested whether pools could be used to elucidate chemical-genetic associations. Three antimalarials with established molecular targets were selected for initial testing: the dihydroorotate dehydrogenase (DHODH) inhibitor, DSM1, the phosphatidylinositol 4-kinase (PI4K) inhibitor, KDU691, and the P-type ATP-dependent Na^+^ pump (ATP4) inhibitor, cipargamin (aka KAE609, NITD609)^27,28,29^. The *P. knowlesi* pool previously transfected with 96 PbACs – and containing these three target genes - was separately exposed to each compound at 3xIC_50_ concentration (determined experimentally against *P. knowlesi* WT parasites; Figure S4) over 12 days of continuous culture. Selection with DSM1 rapidly selected for resistant parasites, resulting in continued growth of the mutant pool under selection. In contrast, WT parasite growth was arrested after 48 hours (Figure S5A). Selection with KDU691 and cipargamin did not yield strong resistance phenotypes, with pooled mutants growing only slightly better than WT parasites under KUD691 selection (Figure S5B) and growing at the same rate as WT under cipargamin selection (Figure SC).

Compound selection could result in increased abundance of individual strains regardless of whether resistance phenotypes could be clearly identified in growth assays. To explore this, BarSeq was performed pre- and post-drug selection to establish the fold change of each PbAC barcode over the selection period for each compound. Selection with all three compounds yielded a significant enrichment of their target genes relative to all other barcodes in the pool, indicating that parasites containing the PbACs encoding those genes had a selective advantage under pressure with each drug, causing their expansion in the population at the expense of susceptible strains (Figure 3 & Figure S6). Selection with DSM1 elicited a ∼300-fold increase in the abundance of *dhodh* barcodes (Figure 3A), with no other barcodes being enriched noticeably above background levels. Strikingly, selection with KDU691 precipitated an ∼700-fold enrichment of *pi4k* barcodes post-selection, in addition to a ∼15-fold enrichment of *rab11a* (Figure 3B). While *pi4k* is the known target of KDU691, *rab11a* has been hypothesised to act as an effector protein downstream of *pi4k* activity, and *rab11a* point mutations have previously been demonstrated to confer resistance to KDU691 in *P. falciparum*^28^, though no copy number variation (CNV) relationship between *rab11a* and KDU691 has previously been established. Here, chemogenomics appears to select not just for the target gene, but also for other genes in the same pathway. Cipargamin selection also elicited a clear enrichment of *atp4* in the pool of 96 PbACs (Figure 3C), although not to the same extent as the other target-gene combinations. For each compound, barcodes corresponding to their target gene also represented the most abundant PbAC detected among the pool of 96 by BarSeq post-selection (Figure S6).

**Figure 3.**
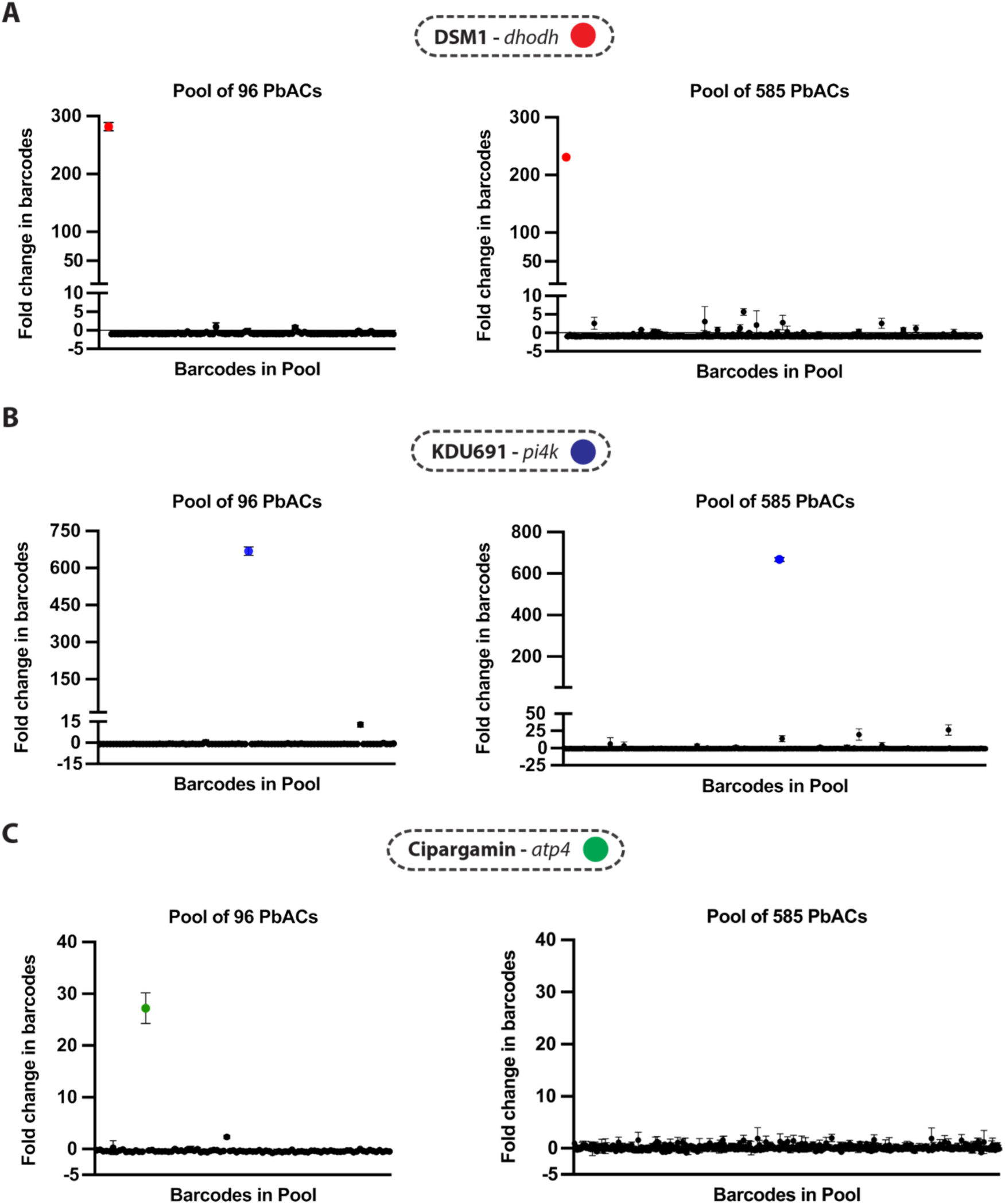
Identifying known chemical-genetic associations using the BarSeq chemogenomics approach. Fold change in barcodes determined by BarSeq following selection of the pool of 96 (left) and 585 (right) PbAC overexpression lines with DSM1 (**A**) KDU691 (**B**) and cipargamin (**C**) at 3xIC_50_ for 12 or 14 days. Each point denotes the proportion of total barcodes represented by a given PbAC barcode in the pool following selection, relative to untreated samples (fold change). Only barcodes which were detectable in the input sample pool at an abundance of > 0.1% (Pool of 96 PbACs) or > 0.015% (Pool of 585 PbACs) and whose fold-change measurement had a 95% confidence interval of < 20 are depicted on graphs. Enriched target barcodes are coloured according to the legend above each panel. Error bars represent SD.

Selection of these antimalarials against a pool of 96 barcoded strains resulted in the clear and unambiguous identification of their target genes, with limited levels of background variation in PbAC abundance. Considering that there are potentially thousands of drug target genes encoded in the *Plasmodium* genome, increasing pool complexity would signficantly increase the throughput of compound screening, and target identification. To test the limits of pool complexity, the entire barcoded PbAC library of 585 PbACs was transfected in pools of roughly identical size, generating six *P. knowlesi* parasite pools each containing ∼100 PbACs. Pools were then combined into a single master pool – theoretically containing the entire barcoded PbAC library – and selected with DSM1, KDU691 and cipargamin at 3xIC_50_ for 14 days (Figure S5D-F). BarSeq identified a clear enrichment of parasites containing the *dhodh* PbAC, demonstrating an increase of over 200-fold post-selection (Figure 3A). Similarly, selection with KDU691 elicited a powerful enrichment of *pi4k* barcodes, increasing over 600-fold (Figure 3B). However, the minor signal for the target-associated gene *rab11a* was lost among this more complex pool (Figure 3B). Both *dhodh* and *pi4k* were also by far the most abundant barcodes detected among the pool post-selection (Figure S6). Enrichment of *atp4* barcodes was not detectable above background following cipargamin selection, suggesting some reduced sensitivity associated with the larger pool (Figure 3C). Nonetheless, the ability of BarSeq to detect significant enrichment of 2/3 target genes in a complex pool of 585 strains after a short selection period indicates the utility of this larger pool for future systematic screening efforts.

### Screening antimalarial phenotypic hits to identify novel targets

Having utilised the PbAC library to identify known gene-drug associations in *P. knowlesi*, we sought to determine if this approach could be used to identify the targets of early-stage antimalarial candidates with unknown mechanisms of action. To this end, a set of 9 inhibitors (Table S2), all with validated activity against *P. falciparum* parasites *in vitro* but with unknown molecular targets, were screened against the entire 585 PbAC library pool. The inhibitory activity of the compounds was first tested against *P. knowlesi,* where all compounds exhibited an IC_50_ in the low micromolar range (Figure 4A, Figure S7 & Table S3). Subsequently, each compound was used to select the combined 585 PbAC pool and subjected to BarSeq analysis. The majority of compounds failed to result in the enrichment of any barcode above background levels (Figure 4B, Figure S8, Figure S9 & Table S4), indicating that no particular PbAC in the pool conferred a selective advantage under selection. However, selection with Compound 18 resulted in an over 800-fold enrichment of *pi4k* barcodes post-selection, strongly indicating that this compound may be an inhibitor of *pi4k.* No other barcodes, including those of *rab11a,* were enriched above background levels.

**Figure 4.**
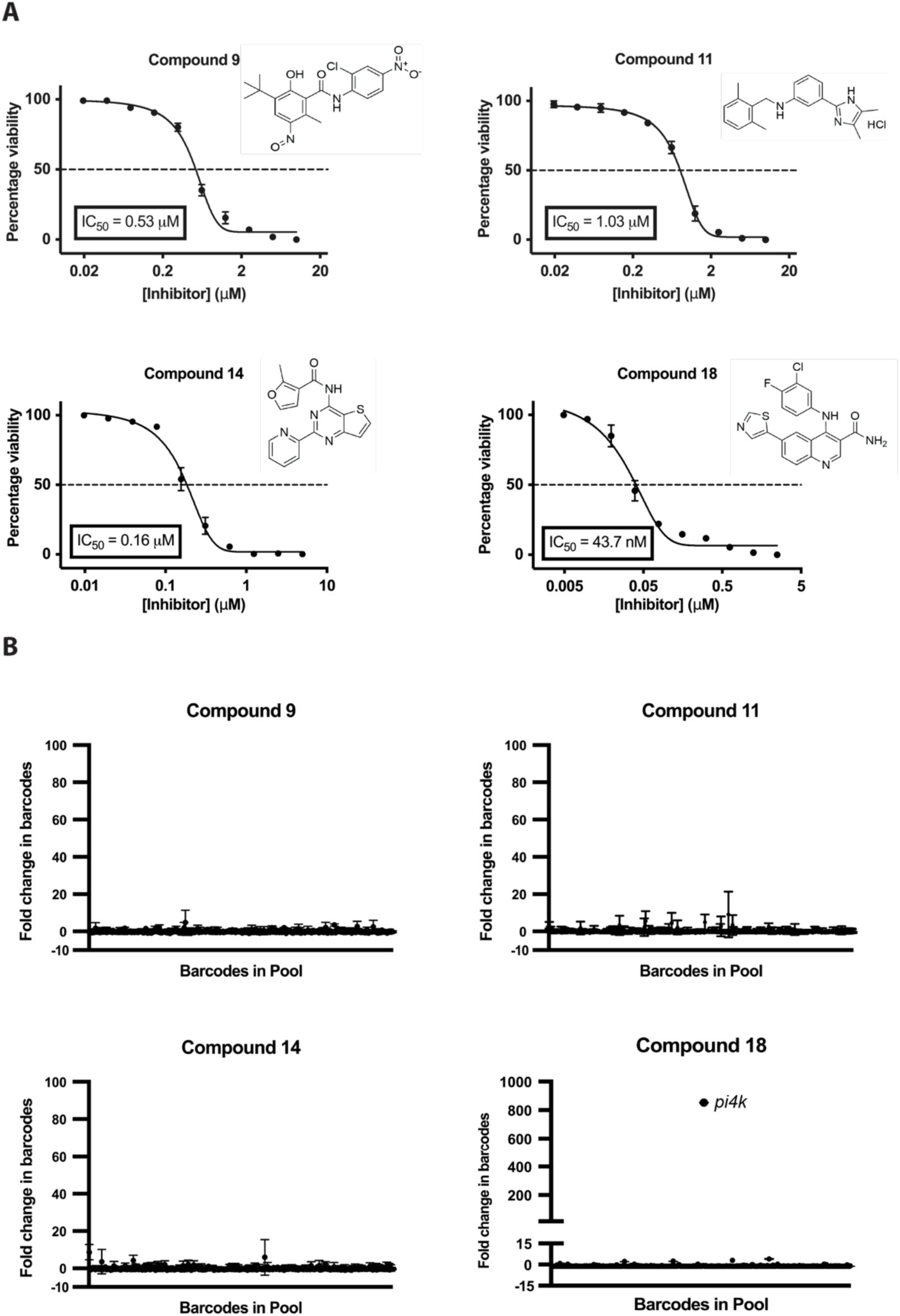
Selecting antimalarial inhibitors with unknown targets using the BarSeq chemogenomics approach. (**A**) Serial dilution dose-response to determine the activity of the GSK compound panel against *P. knowlesi* WT parasites. Error bars represent SD. (**B**) Fold change in barcodes determined by BarSeq following selection of the pool of 585 PbACs with the GSK compound panel. Each point denotes the proportion of total barcodes represented by a given PbAC barcode in the pool following selection, relative to untreated samples (fold change). Only barcodes which were detectable in the input sample pool at an abundance of > 0.015% and whose fold-change measurement had a 95% confidence interval of < 20 were depicted on graphs. Enriched target genes in each case are annotated. Error bars represent SD.

### Validating the Compound 18-*pi4k* association

To validate the Compound 18-*pi4k* association we carried out IVIEWGA in *P. falciparum*. Firstly, the IC_50_ of Compound 18 was validated against *P. falciparum in vitro* (Figure S10), then resistance selection with Compound 18 was performed at 3xIC_50_ using a mutator *P. falciparum* line (Dd2-Polδ) possessing a modification in DNA polymerase δ, resulting in defective proof-reading and a higher propensity to select for resistance^29^. After ∼3 weeks, parasites were recovered in all three wells and clonal lines were isolated. Representative clones (1H5, 2D4, 3C3) from each of the independent selections were evaluated against Compound 18 and exhibited a 6-17 fold shift in IC_50_ (Figure 5A). Consistent with a mode-of-action involving PI4K, the Compound 18-selected clones showed cross-resistance to a known PI4K inhibitor, KDU691^28^ (Figure 5B).

**Figure 5.**
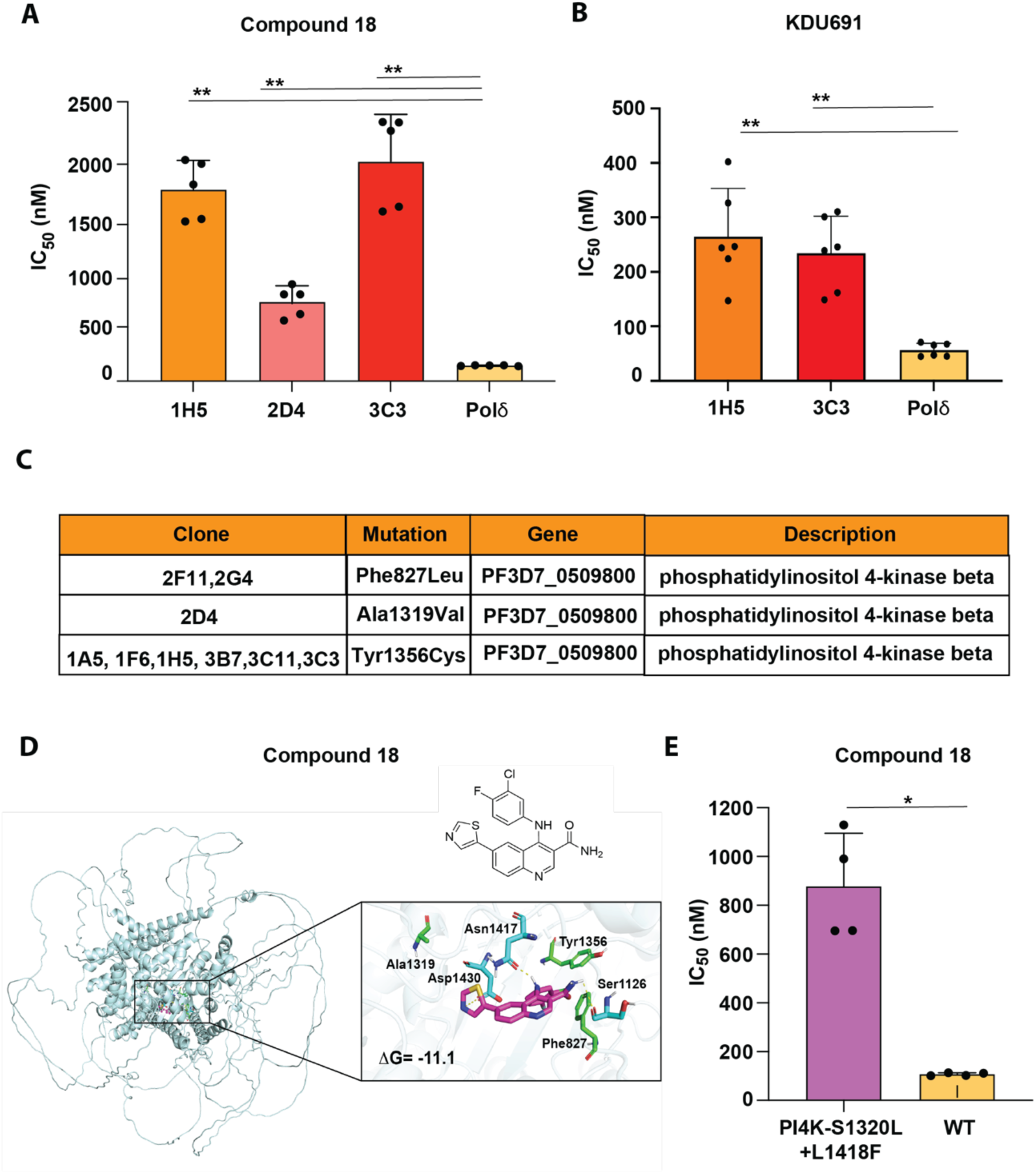
Validation of PI4K as the target for Compound 18. (**A**) Comparison of the IC_50_ values of Compound 18 vs resistant clones generated by *in vitro* evolution using the Dd2-Pol𝛿 mutator line. Clones from three independent flasks were evaluated, alongside the parental Dd2-Pol𝛿 (Pol𝛿) line as a control. (**B**) Assaying clones from two independent Compound 18-selections for cross-resistance to the PI4K inhibitor KDU691. (**C**) Mutations in PI4K identified by whole genome sequencing of resistant clones derived from Compound 18 selection (see Table S5 for full list of single nucleotide variants). (**D**) Virtual docking of GSK Compound 18 into an AlphaFold PI4K protein structure (AF-Q8I406-F1). Active site residues mutated by Compound 18 selection are annotated in green. Residues forming direct interactions with Compound 18 are annotated in cyan. Compound 18 is depicted in magenta. The binding affinity (ΔG) of the Compound 18-PfPI4K interaction is displayed in the bottom left of the docking simulation image. (**E**) IC_50_ values for Compound 18 determined against an independently-derived PI4K mutant and WT control. For all assays, each dot represents a biological replicate (n=4-6) with mean±SD shown as bar chart, and statistical significance determined by Mann-Whitney *U* test (*p<0.05, **p<0.01).

Next, we performed whole genome sequencing (WGS) of nine resistant parasite clones, three each derived from each independent selection. Comparison of the clones to their isogenic parent identified 34 missense mutations in 32 core genes (Table S5). Notably, all clones had mutations in phosphatidylinositol 4-kinase beta (PI4KIIIβ, PF3D7_0509800) (Figure 5C) an enrichment not expected by chance given that there are almost 6000 genes in the genome. F827L and A1319V mutations were present in clones derived from one independent culture, whereas clones from the other cultures possessed the Y1356C mutation, associated with higher-level resistance to Compound 18. Both A1319V and Y1356C localise to the lipid kinase domain, with F827L present on a nearby loop (Figure 5D)^28,30^. Simulated docking of Compound 18 into a predicted PfPI4K structure highlights interactions with several residues close to or located within the lipid kinase domain (S1126, N1417, and D1430), and a binding conformation with close proximity to the residues which confer resistance to Compound 18 when mutated (Figure 5D).

Finally, we tested Compound 18 against an independently-derived *pi4k* mutant parasite (PI4K-S1320L+L1418F)^28^, which demonstrated an ∼8-fold shift in IC_50_ against Compound 18 (Figure 5E). Collectively, the evolution of multiple PI4K mutants under Compound 18 pressure, the cross-resistance of these mutants against known PI4K inhibitors, and the elevated IC_50_ of Compound 18 against an unrelated *pi4k* mutant parasite are consistent with *pi4k* being the target of Compound 18, validating the identification of this target using the PbAC/*P. knowlesi* chemogenomics approach.

## Discussion

This chemogenomic approach represents a novel and unique platform for antimalarial target ID, utilising the systematic overexpression of targets in combination with BarSeq to deconvolute drug mechanisms of action in *Plasmodium* parasites. Data presented here suggests that the approach possesses several notable advantages as a target ID methodology. BarSeq is highly effective at detecting subtle differences in strain sensitivity in pools during drug selection, including those not detectable in conventional growth assays, ie where no clearly resistant parasites emerge. Hits are high-confidence, with target barcodes often enriching several hundred-fold above background levels - consistent across a range of pool complexities – and a low false positive rate, which reduces the likelihood that precious resources are deployed to follow up false leads. The enrichment of *rab11a* in addition to *pi4k* following selection with KDU691 suggests that non-target resistance-associated genes may be identified, providing additional information about target pathways beyond the target of action itself. Only brief durations of compound selection (between 12-14 days) are required to enrich targets to high levels, with the short life cycle of *P. knowlesi* (∼27 hours *in vitro*), allowing resistant parasites to multiply and expand in pools rapidly. This platform is also inherently scalable; compounds can be screened against >500 targets in a single well, with minimal amounts of parasite material required to amplify barcodes for BarSeq. Moreover, all stages of the target ID workflow (parasite culture, drug selection and NGS) are amenable to assay miniaturisation and automation, meaning that the approach could be scaled up to screen compounds and identify targets in much higher throughput than employed here.

PI4K represents a robustly characterised druggable target in *Plasmodium*, with extensive chemical validation, including as the molecular target for imidazopyrazine, quinoxaline^28^, 2-aminopyrimidine^30^, napthyridine^31^, and bipyridine-sulfonamide^32^ chemical classes, in addition to the structurally-distinct Compound 18. Several PI4K inhibitors have progressed to pre-clinical assessment, demonstrating high selectivity for *Plasmodium* PI4K over the nearest human orthologue (PI4KIIIβ), owing to substantial sequence, structure and domain architecture differences between the two kinases^33^, while the most advanced PI4K inhibitor, MMV390048^30^, has progressed to clinical trials^34,35^. It is unclear why many diverse chemical scaffolds converge upon PI4K, though this may reflect the relative paucity of druggable targets in *Plasmodium* in addition to the vital biological role performed by this protein in the parasite. Attempts to introduce a stop codon mutation into the catalytic domain of *Pfpi4k* demonstrate its essentiality for *P. falciparum* growth in the asexual blood stages^28^ while chemical inhibition of PI4K blocks transmission and liver stage development^30^. PI4K is a type IIIβ phosphatidylinositide kinase responsible for the generation of PI(4)P from phosphatidylinositol (PI), which recruits effector proteins at the Golgi to generate transport vesicles, a vital function in the regulation of intracellular membrane trafficking^36^. Disruption of the intracellular distribution of PI(4)P perturbs effector recruitment and lipid transport, impacting the biogenesis of merozoites during schizogony^28^. PI4K therefore represents a biologically and chemically validated target for antimalarial inhibition.

As with all existing antimalarial target ID methodologies, this approach has limitations. While false positives appear rare due to the high signal to noise ratio, potential false negatives appear more common, with only 1/9 inhibitors from the blinded screen yielding a hit. It is possible that these compounds targets are not present in the current PbAC pools, which enompass only a subset of the *P. berghei* genome, omitting many potential drug targets. However, there exists the capacity to expand the library to include many of the ∼2000 potentially druggable *Plasmodium* genes^25, 37^, via the addition of barcodes to further PbAC constructs. In other cases, meaningful target overexpression may not be achieved, possibly due to incomplete transcription, translation, folding or localisation of *P. berghei* orthologues in *P. knowlesi*, or low PbAC copy numbers driving insufficient target overexpression. Though the expression of several *P. berghei* genes has been validated by RT-qPCR, and the generation of extra target protein essentially confirmed by BarSeq for DHODH, PI4K and ATP4, we cannot yet estimate what proportion of the library is successfully overexpressed. Expressing genes from one *Plasmodium* species in another is used relatively widely in the field but it may not always be successful. However, chemical-genetic associations identified using this approach should be conserved across *Plasmodium* species, as both *P. berghei* and *P. knowlesi* are sensitive to most known antimalarial agents, which largely act via the same mechanisms of action across *Plasmodium* species^38^. Other factors accounting for false negatives could include compounds not directly inhibiting enzymatic targets (e.g. Chloroquine/4-aminoquinolines), or hitting multiple targets, differential sensitivity of target orthologues to compounds between different *Plasmodium* species, targets constituting multi-subunit complexes and those interacting with host targets.

These limitations aside, this chemogenomic platform represents a novel and useful addition to the compendium of target ID approaches in *Plasmodium*. The key attribute of scalability means that hundreds of compound-target interactions can be screened in a single culture plate, and the low false positivity rate means that there is minimal risk that significant and precious resource will be invested into following up false leads. Carrying out PbAC chemogenomic screens at an early stage of a target identification pipeline, potentially in parallel with *in vitro* genome evolution, has the potential to signficantly speed up the identification of compound-gene interactions, and hence speed up the development of much-needed new antimalarials.

## Acknowledgements

Funding: This work was supported by the Wellcome Trust [JR: 220266/Z/20/Z, Sanger: 206194/Z/17/Z], the University of Cambridge Wellcome Institutional Partnership Award Developing Concept Fund, the Tres Cantos Open Lab Foundation, project TC266. M.R.L. was supported in part by a Ruth L. Kirschstein Institutional National Research Award from the National Institute for General Medical Sciences (T32 GM008666). This publication includes data generated at the UC San Diego IGM Genomics Center utilizing an Illumina NovaSeq 6000 that was purchased with funding from a National Institutes of Health SIG grant (#S10 OD026929). We are grateful to the staff in the Sanger Scientific Operations for their support with sequencing.

## Author Contributions

Conceptualization, J.C.R. and C.B-L.; Methodology, C.B-L. and J.C.R., Software, F.S. and R.C.; Investigation, C.B-L., M.R., W.C.K., G.G., M.R.L., K.C..; Writing – Original Draft, C.B-L., and J.C.R.; Writing – Review & Editing, C.B-L., J.C.R., S.M-C., M.G.G-L., F-J.G., E.A.W., O.B., M.C.S.L.; Funding Acquisition, J.C.R., M.C.S.L., E.A.W.; Resources, M.C.S.L., E.B., O.B., S.M-C., M.G.G-L., F-J.G.; Visualisation, C.B-L., M.R. K.C.; Supervision, J.C.R., M.C.S.L., O.B., E.A.W., S.M-C., M.G.G-L., F-J.G.

## Supplemental Information

Document S1. Figures S1-S10

Table S1. List of constructs in the barcoded PbAC library.

Table S2. Chemical structures for the GSK inhibitor panel.

Table S3. IC50 values for compounds screened in this study determined against *P. knowlesi*.

Table S4. Barcode counts for the pool of 585 PbACs selected with the GSK inhibitor series.

Table S5. List of SNVs identified following selection of P. falciparum with GSK Compound 18.

Table S6. List of oligonucleotide primers used in this study.

## STAR Methods

### Selection of PbAC library for barcoding

PbAC constructs selected for the addition of a barcode for chemogenomic screening were those fulfilling all of the following three criteria; i) Genes which corresponded to a PbAC in the library containing a promoter region of ≥ 1kb and a terminator region of ≥ 0.5 kb either side of the ORF of interest (determined using data available from https://plasmogem.umu.se/pbgem/); ii) Genes which were annotated as enzymes, transporters, channels or membrane proteins (identified using the search function from https://plasmodb.org/plasmo/app); iii) Genes expressed in the liver stages of infection, identified based on available transcriptomic data sets^39,40^, after which genes which were previously experimentally determined to be redundant in *P. berghei* liver stage development were then removed^41^. This culminated in 730 candidate PbACs for barcoding. Although prioritised based primarily on liver stage expression and non-redudancy, there is significant overlap in *Plasmodium* gene expression between erythrocytic and exo-erythrocytic life-cycle stages^42^, with ∼85% of the 730 candidate PbACs also constituting genes expressed and non-dispensible in the bloodstream stages of infection. It was reasoned that these candidates could serve as a resource for chemogenomic experiments in both liver and bloodstream stages, with the small minority of genes not occupying these criteria in each case serving as negative controls in target identification assays.

### Recombineering PbACs with barcodes

PbAC library constructs were obtained from *Plasmo*GEM (https://plasmogem.umu.se/pbgem/) via the Wellcome Sanger Institute. Individual PbAC constructs expressed by *E. coli* (BigEasy TSA electrocompetent cells, Lucigen) were stored as frozen glycerol stocks and inoculated into 1x Terrific Broth (TB, Invitrogen) supplemented with 0.4% v/v Glycerol, 12.5 µg/µl chloramphenicol and 0.2 mM IPTG in 1 mL volume 96-well deep-well plates and grown at 37 °C for 16 hours. Individual clones were pooled (up to a maximum of 96 at a time) and diluted in TB to an OD_600_ of 0.05, then incubated shaking (220 rpm) at 37 °C until reaching an OD_600_ of 0.6-0.8. Pooled cultures were chilled on ice for 15 min before 1.4 mL of culture was pelleted (5000 rcf, 3 min, 4 °C) and washed 3 times in ice cold water. Pellets were resuspended in 1 mL water and transferred to a 1 mm electroporation cuvette prior to transfection (1800 V, 25 µF, 200 Ω) with a plasmid containing the coding sequences for a 5’ exonuclease in addition to β and γ recombinase enzymes from the lambda phage red operon. Transfectants were recovered in 950 µl TB supplemented with 0.4% glycerol for 70 min at 30 °C, then cultured overnight at 30 °C in TB supplemented with chloramphenicol, IPTG (same concentrations as above) and 5 µg/mL tetracycline. Cultures were diluted to an OD_600_ of 0.05 and incubated shaking at 220 rpm until reaching an OD_600_ of 0.2-0.4, after which expression of the recombinase genes was induced by addition of 10% L-arabinose followed by stationary incubation at 37 °C for 5 min and shaking incubation at 37 °C for 35 min. Cells were pelleted, washed and transfected (as described above) with linear DNA containing regions of homology to the PbAC vector backbone, a zeocin resistance cassette and a randomly generated 12 nucleotide barcode. Cells were recovered and cultured in TB supplemented with chloramphenicol, IPTG and 50 µg/mL zeocin at 37 °C for 16 hours. 200 µl culture was then spread on LB-agar plates containing zeocin, chloramphenicol and IPTG and incubated at 37 °C overnight. Colonies from plates were expanded in TB containing chloramphenicol, IPTG and zeocin prior to plasmid isolation using the Spin Miniprep Kit or Midi Plus Kit (both Qiagen) and verification of the inserted sequences by Sanger sequencing. Barcoded PbACs were subsequently stored as glycerol stocks at −80 °C.

### Preparing PbACs for transfection

Barcoded PbAC clones were inoculated from individual glycerol stocks into 1 mL TB supplemented with chloramphenicol in 1 mL 96-well deep-well growth blocks and incubated shaking at 37 °C for 16 hours. Equal volumes of culture from individual clones were collated (final volume dependent upon desired pool complexity, between 1 and ∼100 PbACs), with the total pre-inoculum volume not exceeding 1/10^th^ of the final culture volume (between 100-500 mL) in 2***×*** low-salt LB, supplemented with chloramphenicol, IPTG, zeocin and 1***×*** L-arabinose (from 1000***×*** stock) then incubated shaking at 37 °C for 16 hours. Cell pellets were collected by centrifugation (3500 rcf, 40 min, 4 °C) and PbAC DNA was extracted using the QIAfilter Plasmid Maxi Kit (Qiagen) according to the manufacturer’s instructions. Prior to transfection, 30 µg PbAC DNA was digested overnight at 37 °C using NotI-HF (NEB) in a 150 µL reaction following standard NEB restriction digest protocols, in order to free the PbAC telomere arms for elongation by endogenous cellular telomerase activity in the parasite. Successful digestion was confirmed by gel electrophoresis on 1% agarose relative to undigested controls.

### Culture of Plasmodium spp

*Plasmodium* culture was carried out in sterile microbiological safety cabinets in a derogated containment level 3 laboratory. Both *P. knowlesi* and *P. falciparum* were grown in leukocyte-depleted, screened donor blood (purchased from NHS Blood and Transplant, Cambridge, UK – ethical approval from NHS Cambridge South Research Ethics Committee, 20/EE/0100, and the University of Cambridge Human Biology Research Ethics Committee, HBREC.2019.40), within canted-neck vented cell culture flasks or multi-well plates in humidified 37 °C incubators supplied with a gas mixture of 3% carbon dioxide and 1% oxygen in nitrogen. Media for *Plasmodium* culture consisted of 500 mL unmodified RPMI 1640 media (Gibco), supplemented with 5 mg/ml Albumax II, 2 mg/ml dextrose anhydrous, 5.96 mg/ml HEPES, 0.3 mg/ml sodium bicarbonate, 0.05 mg/ml hypoxanthine, 1% v/v L-glutamine and 0.1% v/v gentamycin sulfate. Growth media for *P. knowlesi* was additionally supplemented with horse serum at 10% v/v. *P. knowlesi* was cultured at 2% and *P. falciparum* at 4% haematocrit respectively, unless stated otherwise.

In preparation for use in *Plasmodium* culture, blood was washed once in an equal volume of RPMI 1640 media followed by centrifugation (2000 rcf, 5 min, 20 °C), and resuspended in an equal volume of culture media to prepare a working blood stock at 50% haematocrit, which was stored at 4 °C for no longer than 7 days. Fresh blood was obtained on a weekly basis. All centrifugations to separate iRBCs from assay medium (pelleting) were performed at 1100 rcf, 5 min, 20 °C, acceleration 9, brake 3.

Synchronisation of *P. knowlesi* cultures was performed using Histodenz (Sigma Aldrich) prepared at 27.6% w/v in 10 mM HEPES (adjusted to pH 7.0 and filter sterilised) and diluted to 55% v/v in complete media. Transfections of *P. knowlesi* were performed following established protocols^43^. Clonal populations of *P. knowlesi* and *P. falciparum* parasites were obtained using limiting dilution cloning^44^.

### IC_50_ determination

The half maximal inhibitory concentration (IC_50_) values of antimalarial compounds were determined by screening in serial dilution against mixed stages of either *P. knowlesi* A1H1 or *P. falciparum* 3D7 WT parasites, using SYBR Green I (Invitrogen) as a marker of parasite viability^45^. Compounds acquired in powder form were reconstituted in sterile dimethylsulfoxide (DMSO) and stored at −20 °C. Assay plates containing serially diluted drugs and controls were incubated for 48 hours (*P. knowlesi*) or 72 hours (*P. falciparum)* at 37 °C in gassed humidified incubators. Following RBC lysis^45^plates were mixed by agitation at 800 rpm and incubated for a further 30 min at 37 °C in the dark. Fluorescence measurements were made using a Clariostar plate reader (BMG Labtech) using a 485-nm excitation filter and a 535-nm emission filter.

Treating raw fluorescence values from negative control wells (unparasitized RBCs) as 0% and positive controls (parasited RBCs treated with DMSO vehicle-control) as 100%, each drug-treated well was assigned a percentage cell viability value. IC_50_ concentrations were determined by plotting normalised percentage viability (Y-axis) against log-transformed compound concentrations (X-axis) for each compound, then subjected to dose-response non-linear regression analysis. IC_50_ values were determined from data obtained from at least three experimental plates, where each compound was assayed in triplicate wells, and analysis was performed using Graphpad PRISM version 9.

### *P. knowlesi* PbAC drug selection assays

For the longitudinal selection of compounds, cultures were diluted to 1% parasitemia, 2% haematocrit and deposited into 24-well culture plates (Falcon) at a final volume of 1 mL per well. Plates were incubated at 37 °C until the RBCs had sedimented to the bottom of the well, after which the culture supernatant was replaced with complete media containing compounds diluted to desired concentrations, then blood resuspended completely. Each cell line was selected with each compound/concentration in at least duplicate wells. Plates were incubated continuously at 37 °C, and every 48 hours samples were taken for parasitemia determination using flow cytometry (see below). The parasitemia of each well was then adjusted to 1% by dilution with complete media and RBCs if measuring higher than 1%. Plates were incubated further until RBCs had sedimented, then the media was replaced with fresh compound-containing media and subsequently incubated at 37 °C. This process was repeated for between 10-14 days (depending on pool complexity), after which cultures were pelleted, washed once in phosphate buffered saline (PBS) and iRBCs were stored at −20 °C until BarSeq library preparation.

### Flow Cytometry to determine parasitemia

The parasitemia of drug-treated cultures was determined by dispensing 50 µL of PBS into the wells of round bottom 96-well plates (Falcon), to which 20 µL of resuspended culture containing iRBCs was added. At least one well containing uninfected RBCs at the same haematocrit as culture samples was used as a negative fluorescence control. A further 150 µL of SYBR Green I (Invitrogen) diluted in PBS to a final concentration of 2***×*** was added to each well. Plates were covered and incubated at 37 °C for 45 min in the dark, then centrifuged (450 g, 3 min, 20 °C), the supernatant removed, and each blood pellet resuspended in 200 µL PBS. Plates were examined with a 488 nm blue laser to excite SYBR Green I using an Attune NxT Flow Cytometer (Thermo Fisher Scientific) with an attached 96-well plate autosampler (CytKick). 100,000 events per well were measured at a flow rate of 25 µL/min from an acquisition volume of 75 µL to determine parasitemia.

### BarSeq

To prepare BarSeq libraries, the region of the PbAC construct containing the barcode module was amplified by PCR from either 50 ng gDNA extracted from PbAC transfectants using a DNeasy Blood & Tissue Kit (Qiagen) according to the manufacturer’s instructions or directly from 1.5 µL iRBCs using the primers BA2-int1 and BC_STM_91 under the following cycling conditions 98 °C, 5 min // 35 cycles of 98 °C, 30 s / 57 °C, 30 s / 68 °C , 10 s // 68 °C, 5 min // 4 °C hold). Amplified PbAC barcode regions (∼250bp) were resolved on 2% agarose gels with reference to the 100 bp DNA ladder (NEB).

Amplicons were ligated with adapter and index sequences for Illumina sequencing in a single nested PCR reaction using two common PAGE-purified Ultramers and a set of 96 uniquely barcoded indexing Ultramers (both IDT). All primers used in BarSeq library preparation are shown in Table S6. Each primer was used at a final concentration of 600 nM and the PCR template for amplification consisted of 4 µL of each barcode amplicon generated in the previous PCR reaction, (cycling conditions; 98 °C, 3 min // 8 cycles of; 98 °C, 30 s / 55 °C, 30 s / 72 °C for 30 s // 72 °C, 5 min // 4 °C hold). Amplicons produced a band at ∼345 bp when resolved on 2% agarose gels with reference to the 100 bp DNA ladder (NEB). All PCR reactions in the preparation of BarSeq libraries were performed in a final reaction volume of 25 µL using CloneAmp Hifi PCR Premix (Takara Bio) according to the manufacturer’s instructions. All PCR reactions were performed using a C1000 Touch thermal cycler (Biorad).

Indexed samples were quantified by QuBit and pooled at equimolar amounts prior to purification using Agencourt AmpureXP beads (Beckman Coulter™) and a magnetic separation stand according to the manufacturers instructions. The final elution was performed using 300 µL of buffer EB (QIAGEN) overnight at 30 °C.

Pooled libraries were quantified using QuBit and diluted to 8 nM in nuclease-free water prior to sequencing on an Illumina Miseq using a MiSeq Reagent Kit v2 (300 cycle) with the following run parameters; sequence at low cluster density (400 k/mm^2^), 150 bp paired-end reads, pass conditions (% PF ≥ 85) and 50% PhiX spiked in. In order to count PbAC barcodes in each sample, FastQ files were collated into a single directory and barcodes were extracted from each file using a PERL script which selected reads with the correct flanking sequences and counted exact matches of unique barcodes between these constant regions to generate spreadsheets containing the counts for each PbAC barcode in each library sample. PbAC barcodes were transformed into gene IDs corresponding to genes of interest within the gDNA insert of the PbAC. The abundance of each PbAC barcode was calculated in each sample by dividing the number of counts for a barcode by the total number of barcodes in the sample. Fold change in barcode abundance was determined by subtracting the number of counts for each barcode in treated conditions by the number of counts in untreated conditions, then dividing by the number of counts in untreated conditions (Treated – Untreated / Untreated). Thresholds were applied to eliminate barcodes from subsequent analysis with a low abundance in the input transfection pool pre-selection^25^ (0.1% for pools of ∼100 PbACs, 0.015% for pools of ∼600 PbACs) and when plotting fold change data, only barcodes detectable above these thresholds were shown. Similarly, fold change values with unusually high variance (a 95% confidence interval of > 20) were removed, to eliminate barcodes with inconsistent fold change values across replicates. All data analysis and visualisation of BarSeq data was performed in GraphPad PRISM version 9. All sequencing was performed using the Wellcome Sanger Institute Scientific Operations sequencing service.

### RT-qPCR

To measure transcript levels of target genes, late-stage parasites from *P. knowlesi* WT and PbAC transgenic lines were enriched using Histodenz and iRBCs were lysed by resuspension in 10***×*** volume of ice-cold 0.1% Saponin (Sigma) in PBS, and incubated on ice for 10 min. Samples were pelleted, supernatant removed and lysis repeated twice more or until there was no evidence of haemolysis in the supernatant post-centrifugation, retaining a pellet of purified parasite material. Pellets were resuspended in 200 µL PBS with the addition of 300 µL TRIzol (Thermo), incubated at room temperature for 5 min and stored at −80 °C prior to RNA extraction.

Pellets were thawed on ice and resuspended in a further 500 µL of TRIzol, then RNA was extracted using the Direct-Zol RNA Miniprep Plus kit (Zymo Research) according to the manufacturer’s instructions. Contaminating DNA was removed from samples using the Turbo DNase-free kit (Thermo Fisher) and purified RNA was kept at −80 °C for long term storage or on ice and used immediately for cDNA synthesis, which was performed using a High-Capacity Reverse-Transcription Kit (Applied Biosystems). All RNA samples were adjusted to the same concentration prior to reverse transcription to ensure equal reaction efficiencies during cDNA synthesis. cDNA was subsequently stored at −20 °C until use in qPCR experiments.

Prior to gene quantification, the optimal melting temperature (T_m_) for each primer set was determined by assaying primers against positive control templates (purified, amplified PCR products, synthetic gene fragments or pooled cDNA samples) across a thermal gradient, where the annealing temperature producing the lowest C_t_ value for each primer set was used in all subsequent assays. Primers were assayed against pooled positive control templates (cDNA) and post-amplification melt curve analysis was applied to determine the specificity of target amplification. Only primers corresponding to a single melt curve peak were used for gene quantification. Standard curve analysis was used to determine the reaction efficiency of each primer set, where primers were assayed against positive control templates across an 8-point, 10-fold dilution series and C_t_ values (Y-axis) were plotted against log-transformed template dilutions (X-axis). Only primer sets which demonstrated a reaction efficiency of between 90-110% were used to determine gene copy number. All qPCR optimization experiments were performed using PowerUp SYBR Green Master Mix (Applied Biosystems) in a final reaction volume of 10 µL following the manufacturer’s instructions and recommended qPCR cycling conditions. A Bio-Rad CFX96 Real-Time PCR Detection System was used to run all qPCR assays, which were performed using Hard-Shell thin wall PCR Plates (Bio-Rad) and analysed using CFX Maestro software (Biorad).

Target gene quantification was performed in triplicate wells in plates, across at least three experimental plates. To ensure that amplification was not the result of remnant PbAC DNA in cDNA samples, primers were assayed in triplicate against templates subjected to the same reverse transcription protocol as used to generate cDNA from RNA, but without the addition of reverse transcriptase enzyme (-RT). For all primers tested, Ct values recorded in -RT control well were at least 9 cycles greater than those recorded in the corresponding +RT wells (or no Ct value was recorded at all in the -RT sample), indicative of no substantial gDNA contamination in cDNA samples. Primers were also assayed against no cDNA-containing templates, also in triplicate. All analysis, graph plotting and statistical tests were performed in Graphpad Prism. All qPCR primers conformed to a length of 18-30 bp, GC content of 40-60%, mean primer melting temperature (T_m_) of 60 °C (T_m_ of each primer pair not differing by more than 2 °C), minimal runs of identical nucleotides and the presence at least one 3’ GC-clamp. All amplicons were between 75-200 bp in length with a GC content of 40-60%, ideally targeting the 3’ end of transcripts to validate that the entire gene sequence was transcribed. Primers to amplify gene orthologues from different *Plasmodium* species were designed to target regions of the lowest sequence identity to prevent cross-amplification, and where possible, to span introns in order to amplify *P. berghei* cDNA specifically and not gDNA. Each primer set was assayed for secondary structure and self-dimerization using primer analysis software (IDT). Primers were also blasted against the *P. knowlesi* genome using PlasmoDB to eliminate those with significant homology to off-target genomic sequences. All qPCR primers used in these assays are shown in Table S6.

### GSK Inhibitor Panel

All inhibitors used in the blinded compound screen constituted a subset of the Tres Cantos Antimalarial Compound Set (TCAMS) identified as part of high-throughput screens against bloodstream stage *P. falciparum* with an IC_50_ of < 1 µM^7^. All compounds were further validated for dual-stage activity activity against both bloodstream and liver-stages, and confirmed to not act via established mechanism of action pathways, including folate biosynthesis, cytochrome bc_1_ complex inhibition or the electron transport chain^46^ or demonstrating cross-resistance to transgenic lines with mutations in *Pfpft* (protein farnesyltransferase) or *Pfcarl* (cyclic amine resistance locus).

### Resistance selection of *P. falciparum*

*In vitro* drug resistance was generated against GSK Compound 18. Relevant *P. falciparum* parasite lines with an initial inoculum of 1×10^7^ were continuously exposed to 3x IC_50_ of Compound 18 in three independent flasks. This concentration eliminated parasites to a level undetectable by Giemsa-stained thin smears. Parasites were maintained in drug-free media after day 10. Microscopic examination of Giema-stained thin smears was used to monitor parasite death and recrudescence after drug treatment. After recrudescence, drug susceptibility assays were performed on the bulk culture to confirm a shift in IC_50_ in comparison to the parental line. The resistant parasites were cloned by limiting dilution to obtain genetically homogeneous cultures. Nine clones (three from each flask) were harvested for genomic DNA extraction and whole genome sequencing.

### Whole Genome Sequencing and Analysis

Sequencing libraries were prepared using the Nextera XT kit (Cat. No FC-131-1024, Illumina) with 2ng input gDNA and standard dual indexing, then sequenced on an Illumina NovaSeq 6000 at the UCSD Institute for Genomic Medicine (IGM) Genomics Center. The raw sequencing data generated in this study were submitted to the NCBI Sequence Read Archive database (https://www.ncbi.nlm.nih.gov/sra/) under accession number PRJNA1120440. Fastq files containing raw reads were aligned to the *P. falciparum* 3D7 reference genome (PlasmoDB v13.0) and pre-processed following a previously described GATK-based workflow^47^. SNVs and indels were called with GATK HaplotypeCaller^48^, filtered to retain high-quality variants, and variant consequences were subsequently annotated using SnpEff^49^. Mutations that were considered background (native to the compound-sensitive parent) were removed, leaving a only mutations that had evolved over the course of the compound selection process.

### Virtual docking of GSK Compound 18 into PfPI4K

*Plasmodium falciparum* 3D7 PI4K AlphaFold model (AF-Q8I406-F1) was uploaded onto AutoDockTools, along with GSK Compound 18. Water molecules were removed, and atoms were repaired with polar hydrogens and Kollman charges. Autogrid was used to focus a 60x60x60 angstrom box around the three residues identified to confer resistance to Compound 18. Autodock was performed using 50 genetic algorithm runs, with a population size of 300 and evaluation size of 25 million. The 50 resulting conformations were ordered by decreasing binding energies, and the top model was visualized in PyMol.

