## Supplementary Figures for "A novel chemogenomic screening platform for scalable antimalarial drug target identification"

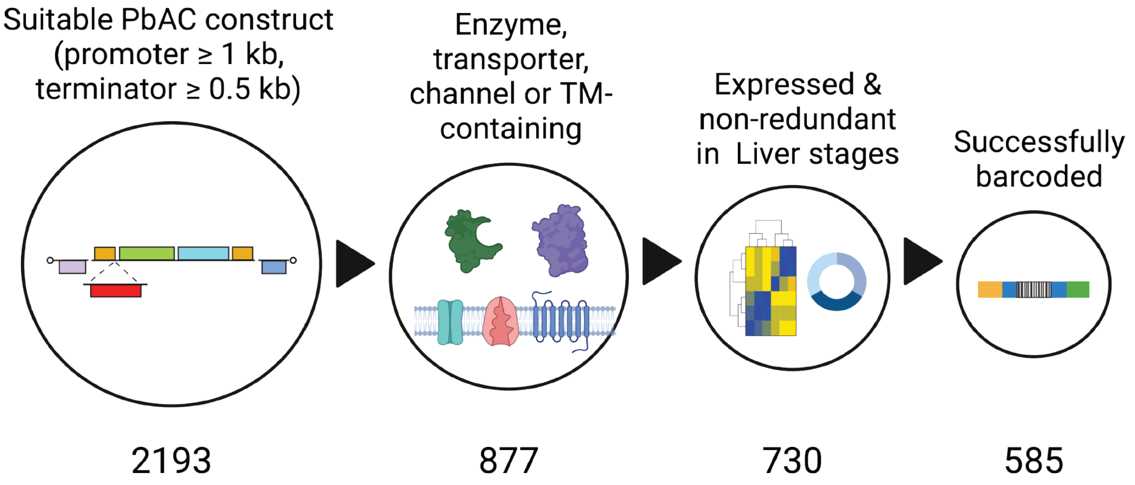


**Figure S1.** Schematic overview of the workflow for the prioritisation of PbACs to be included in the barcoded PbAC library for chemogenomic screening**.** PbACs chosen for barcoding had a promoter region of ≥ 1 kb and a terminator region of ≥ 0.5 kb either side of the ORF of interest to increase the likelihood of successful transcription. Target genes encoded by PbACs were selected on the basis of representing likely drug targets (enzymes, transporters, transmembrane proteins and channels). Genes which were previously experimentally determined to be expressed and non-dispensible in the liver stages of infection were carried forward, and dispensible genes were removed. Barcoding was attempted on a total of 730 PbACs using recombinase-mediated engineering (recombineering), resulting in 585 PbACs which were successfully barcoded.

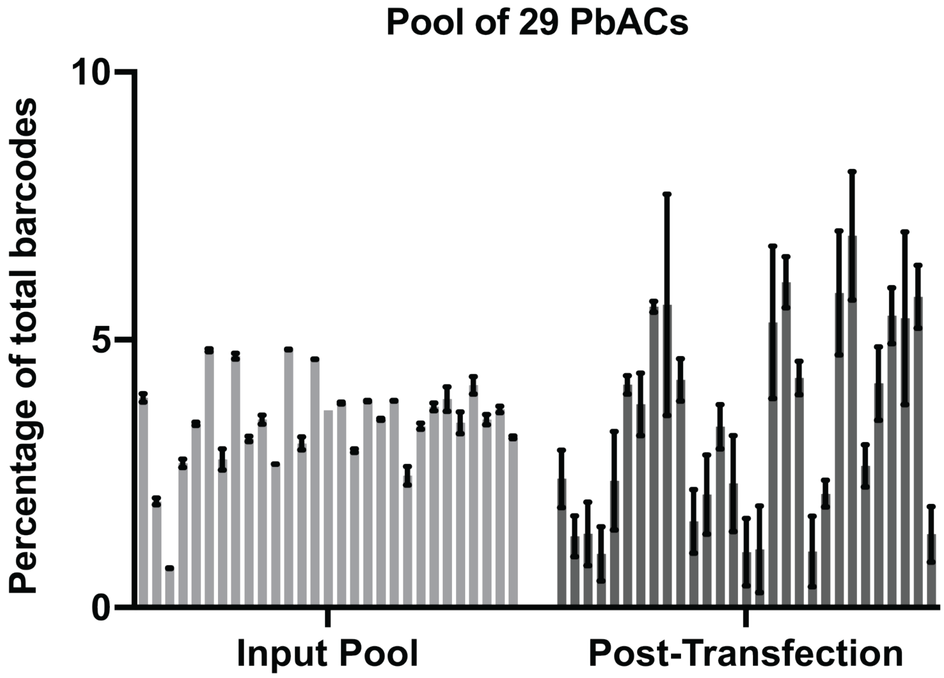


**Figure S2.** Percentage of total barcodes determined by BarSeq for the pre-transfection input pool and post-transfection parasite population transfected with a pool of 29 PbACs. Error bars represent SD, n = 2 replicates for 'Input Pool' and 3 replicates for 'Post-Transfection'.

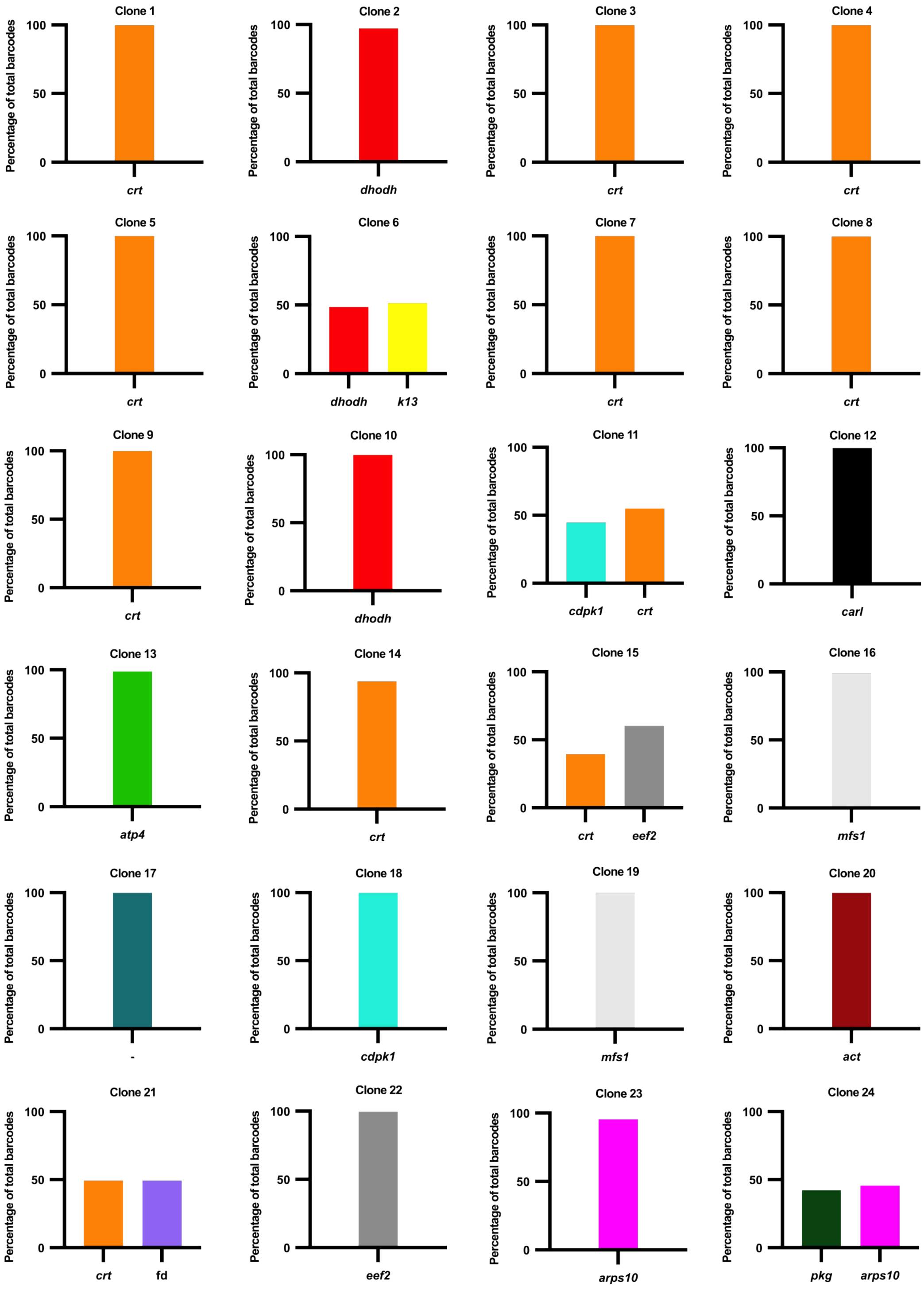


**Figure S3.** Clones obtained from limiting dilution cloning of bulk PbAC transfectant pools. Each clone is represented by an individual graph. Annotated for each clone is the identifying number (1-24) and the gene corresponding to the PbAC barcode(s) detected by BarSeq.

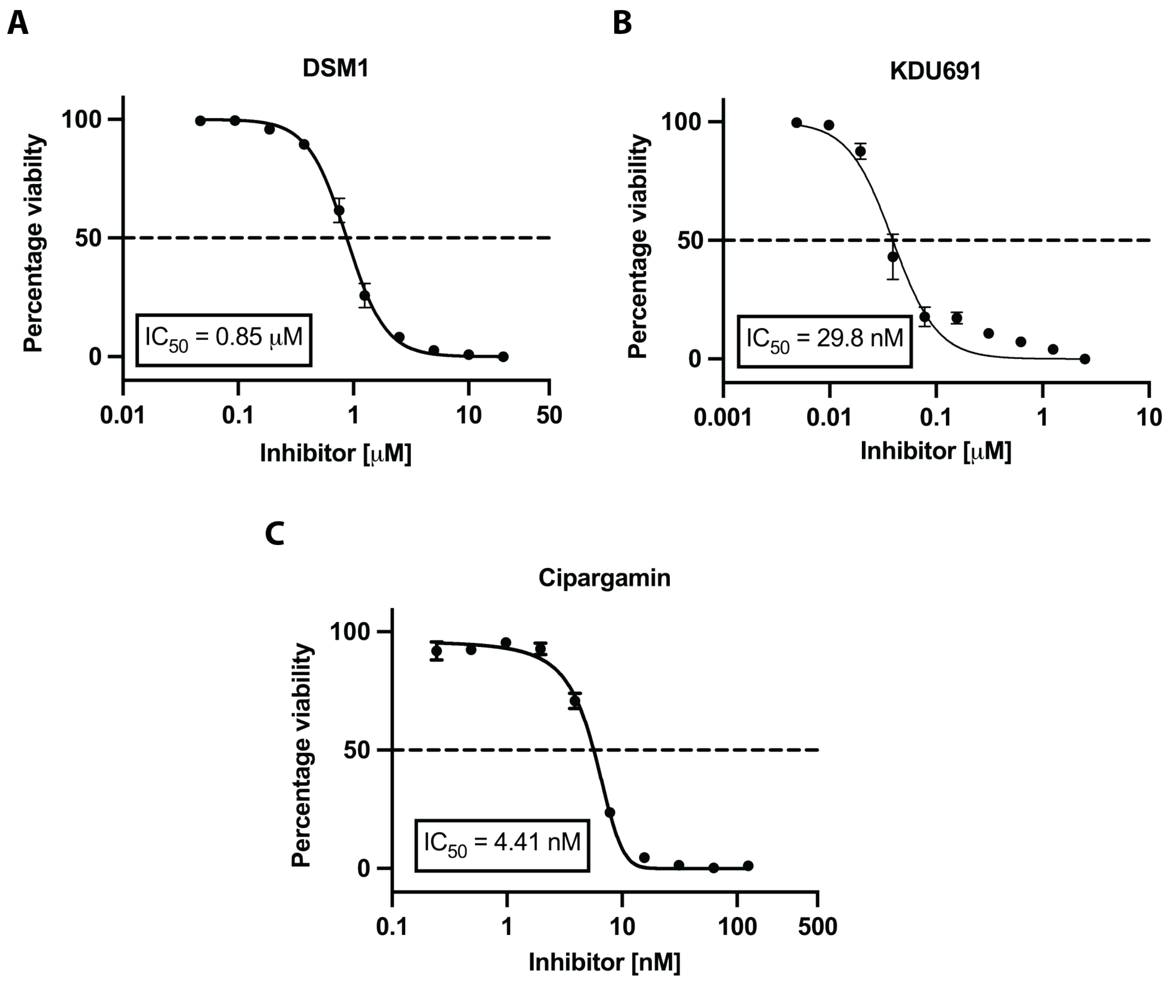


**Figure S4.** Serial dilution dose-response curves for the pilot compounds DSM1 (**A**), KDU691 (**B**) and cipargamin (**C**) assayed against *P. knowlesi* WT parasites to determine their $IC_{50}$. Error bars represent SD. See Supplementary Table S3 for detailed $IC_{50}$ values.

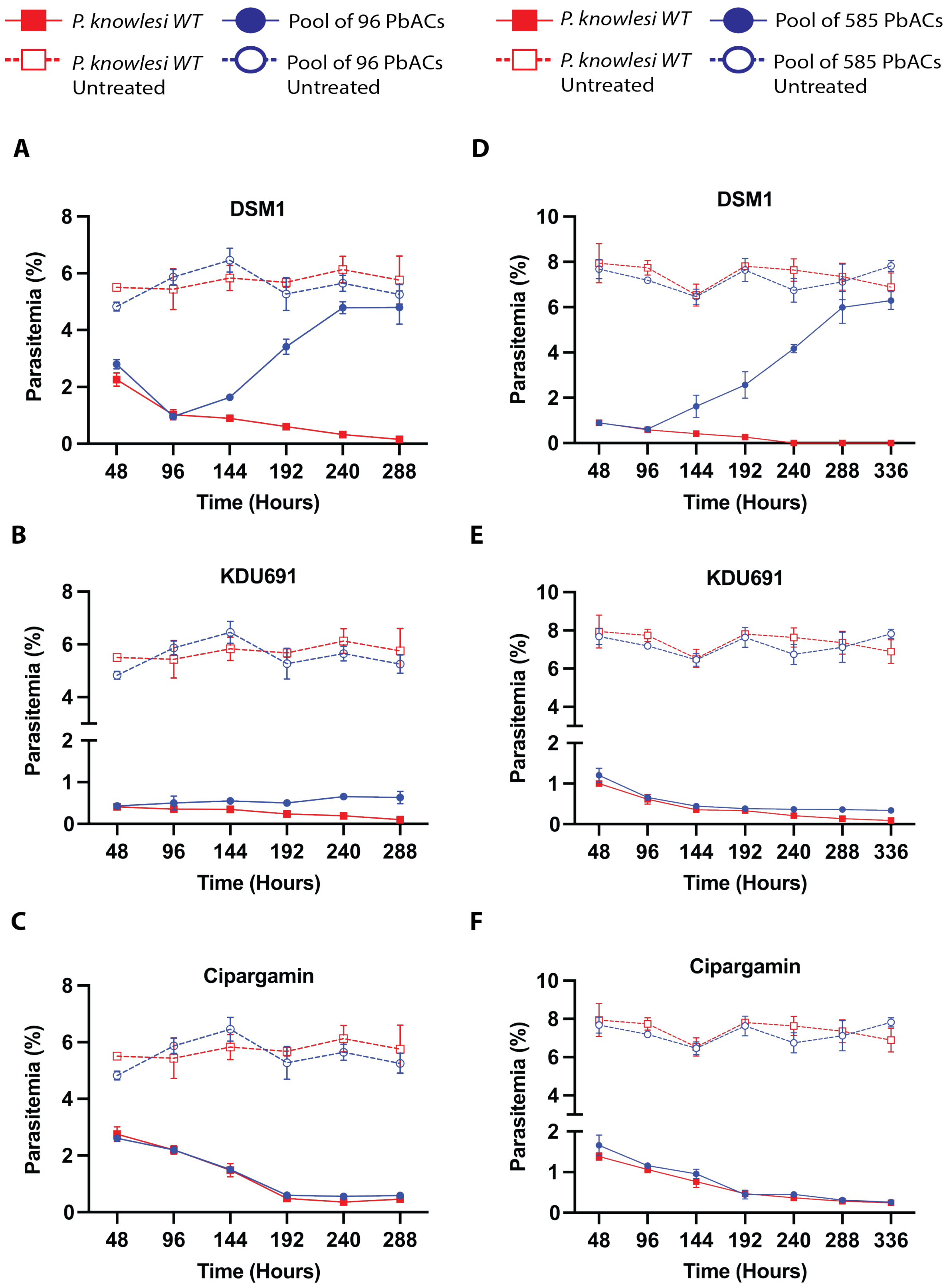


**Figure S5.** (**A-C**) Parasitemia of the pool of 96 PbAC mutant strains and *P. knowlesi* WT parasites selected with pilot compounds DSM1 (**A**), KDU691 (**B**) and cipargamin (**C**). (**D-F**) Parasitemia of the pool of 585 PbAC mutant strains and *P. knowlesi* WT parasites selected with the same compounds. Error bars represent SD.

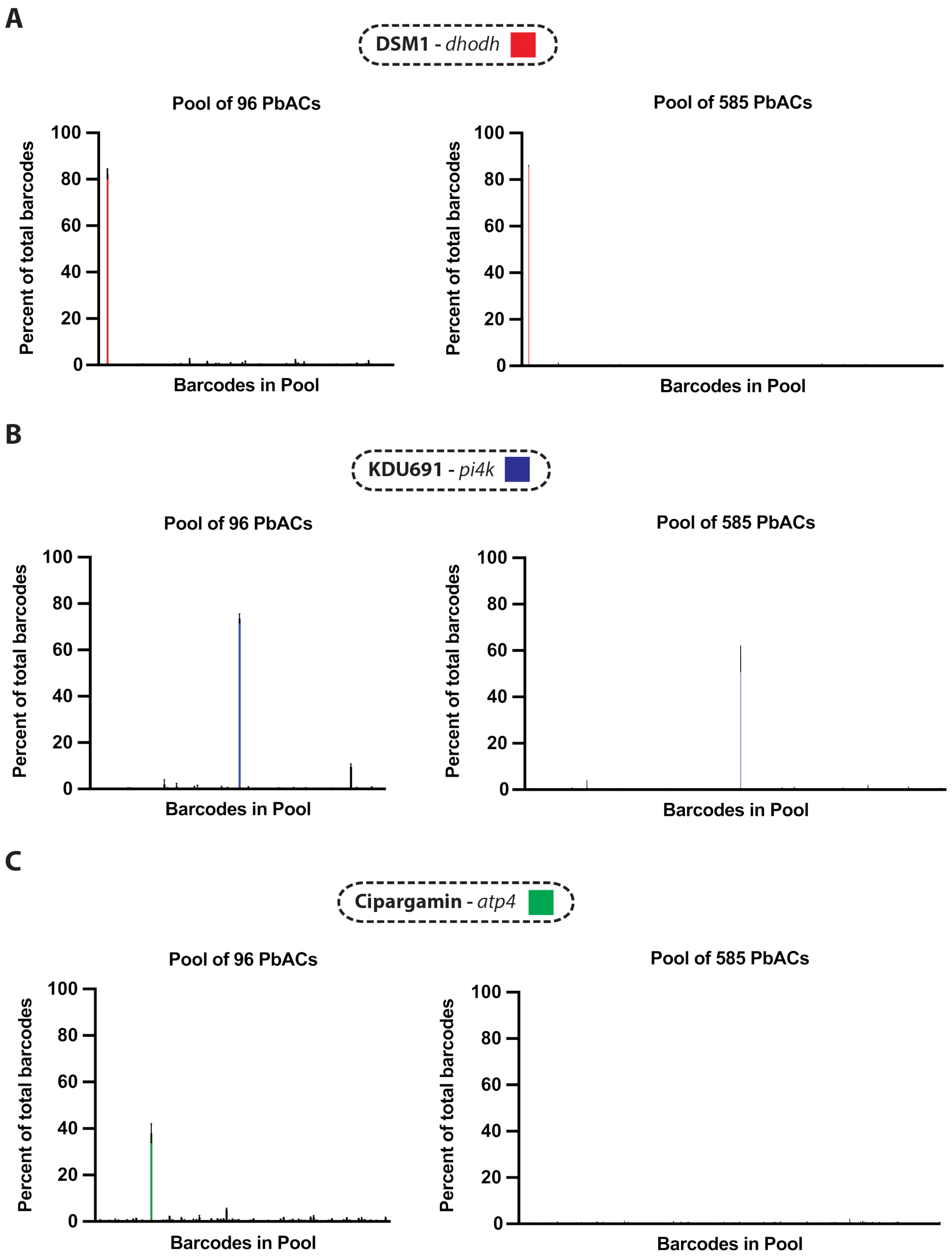


**Figure S6.** Percent of total barcodes determined by BarSeq following selection of the pool of 96 (left) and 585 (right) PbAC overexpression lines with DSM1 (**A**) KDU691 (**B**) and cipargamin (**C**) at 3x $IC_{50}$ for 12 or 14 days. Each bar denotes the proportion of total barcodes represented by a given PbAC barcode in the pool, corresponding to an individual construct in the library. Enriched target barcodes are coloured according to the legend above each panel. Error bars represent SD.

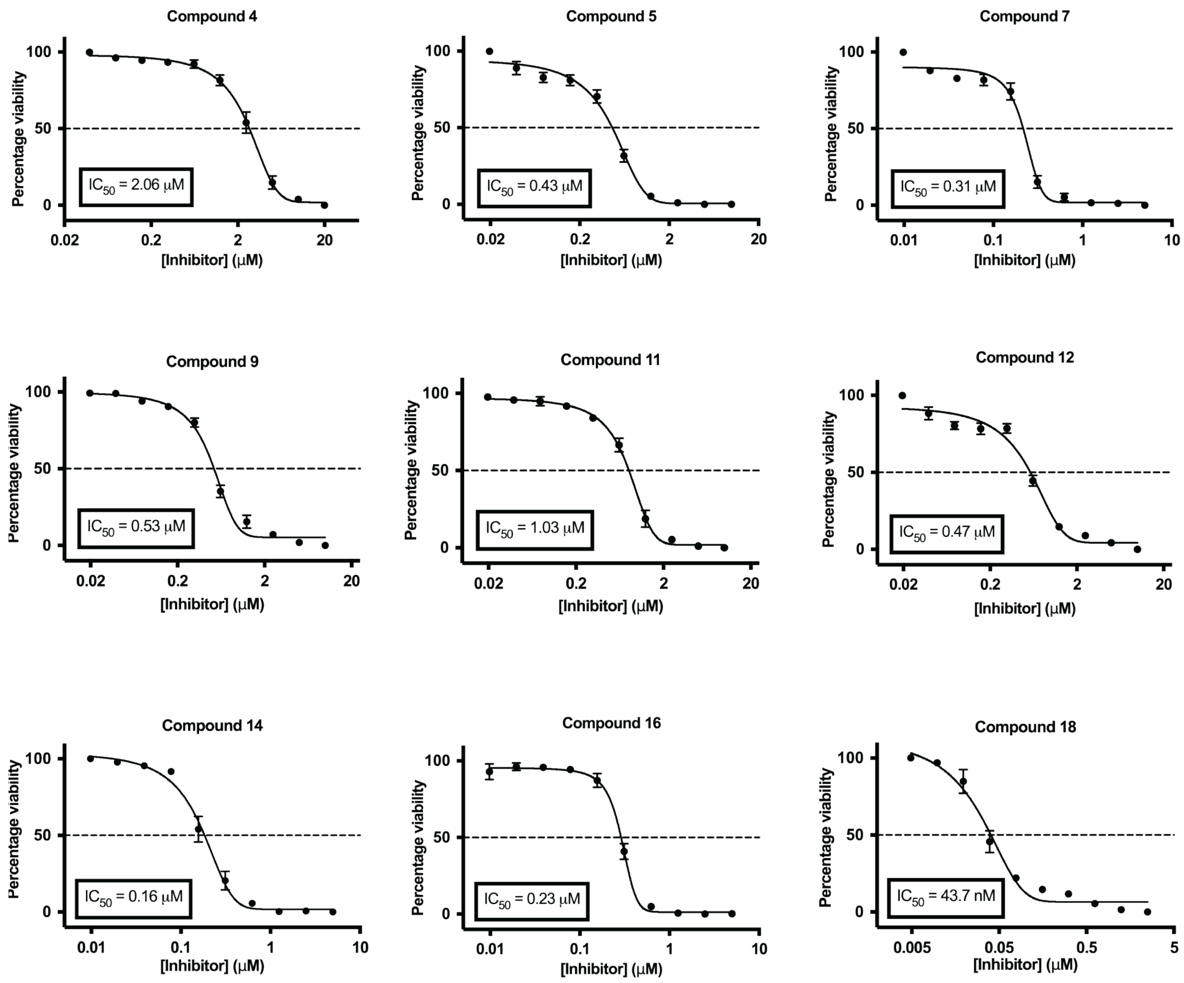


**Figure S7.** Serial dilution dose-response curves for the GSK compound panel assayed against *P. knowlesi* WT parasites to determine their $IC_{50}$. Error bars represent SD. See Supplementary Table S3 for detailed $IC_{50}$ values.

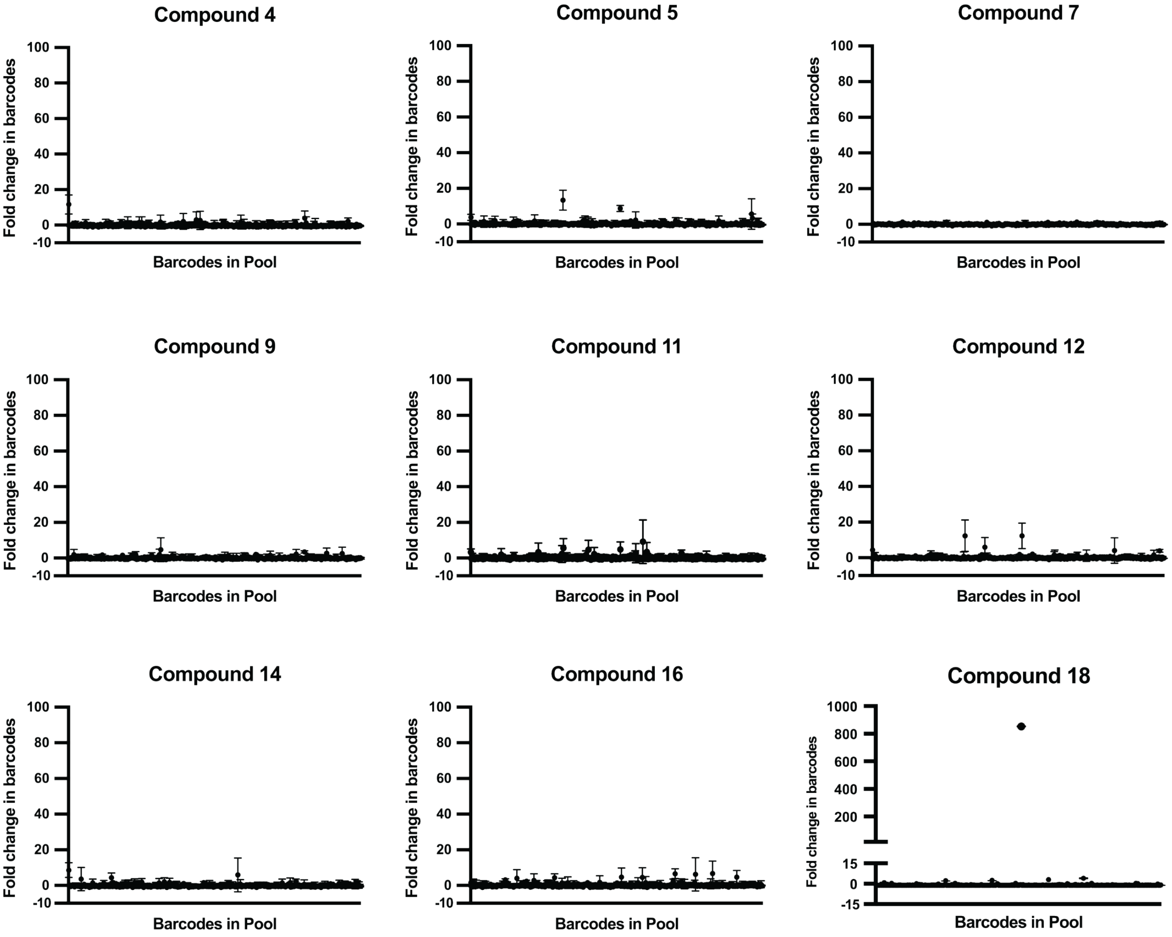


**Figure S8.** Fold change in barcodes pre- and post-selection for the combined pool of 585 PbAC mutant strains selected with the GSK inhibitor panel determined by BarSeq. Each graph represents selection with a different compound from the panel. Only barcodes which were detectable in the input sample pool at an abundance of > 0.015% and whose fold-change measurement had a 95% confidence interval of < 20 were depicted on graphs Error bars represent SD.

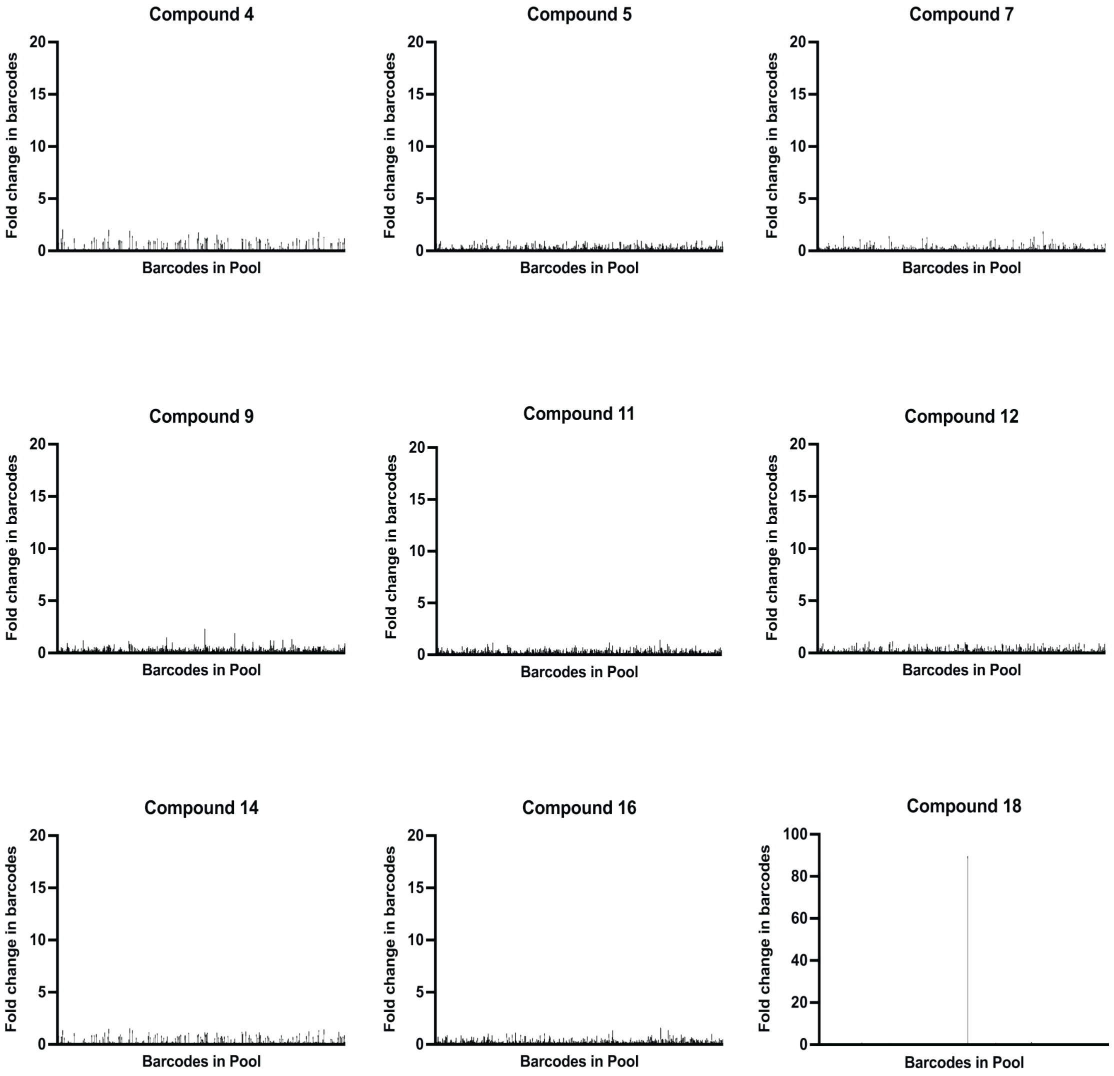


**Figure S9.** Percent of total barcodes determined by BarSeq following selection of the pool of 585 PbAC overexpression lines with the GSK compound panel. Each graph corresponds to a different compound in the panel. Each bar denotes the proportion of total barcodes represented by a given PbAC barcode in the pool, corresponding to an individual construct in the library. Error bars represent SD.

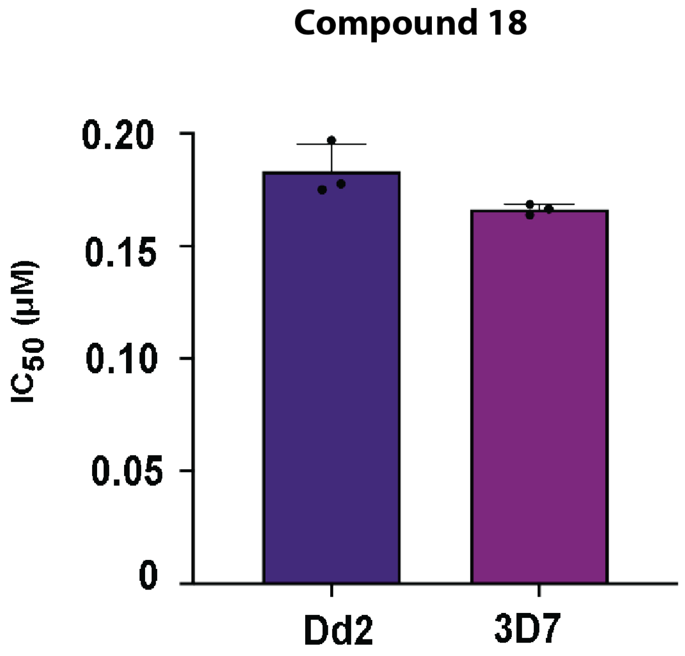


**Figure S10.** $IC_{50}$ determination of Compound 18 against *P. falciparum* Dd2 and 3D7 strains. Each dot represents aa biological replicate (n=3) and error bars represent SD.
